# 3D Multiphoton Nanolithography with Bioresorbable Amino Acid-Based Resins

**DOI:** 10.1101/2025.01.13.632771

**Authors:** Christoph Naderer, Dmitry Sivun, Stefan Haudum, Ian Teasdale, Jaroslaw Jacak

## Abstract

We demonstrate that the newly designed amino acid phosphordiamidate-resins (APdA), containing vinyl reactive groups for polymerization, can be utilized to fabricate sub – 100 nm features through 3D multiphoton lithography. We have quantitatively analyzed the feature size, Young’s modulus, and functionalization of the nanostructures using atomic force and single-molecule fluorescence microscopy. Our results indicate that the polymer backbone, composed of either valine or alanine, imparts hydrophobic properties to the monomer, restricting the swelling of the polymeric nanostructure to 8% in aqueous environments. Despite minimal swelling, experiments revealed an up to 10-fold change of Young’s modulus for dry versus wet conditions. To enhance the versatility of the APdA-based structures, we incorporated biotin functionalization and utilized it for the immobilization of extracellular vesicles. Hence, these findings highlight the potential of APdA-based nanolithography photoresists for biomedicine and nanotechnology applications.

---

3D micro- and nanostructuring with biocompatible materials play a significant role in applications such as tissue engineering, drug delivery systems, biosensors, and microfluidics.^1–6^ For controlled 3D structuring, contemporary techniques such as multiphoton lithography (MPL) allow the fabrication of structures with sub-100 nm feature sizes and sub-micron resolution.^7–9^ In biostructuring, MPL has mainly been linked to synthetic materials,^10^ as seen in 3D structures that support neuronal directional growth^11^ and microcages made from acrylate-based monomers, which have demonstrated increased bacterial mortality.^12^ To enhance the biocompatibility of 3D synthetic scaffolds, various surface modification strategies have been introduced,^13^ including concepts utilizing protein-adhesive photoresists^14,15^ or photoresists with reactive moieties for biomolecule coupling.^16,17^ In terms of nanostructuring, this methodology enables the creation of sub-diffraction-sized features while offering biocompatibility and biofunctionality, both of which are crucial for cell-based 3D scaffold applications. Recent findings on platelet activation have been reported, utilizing individual von Willebrand factor proteins immobilized on a 3D scaffold,^18^ or scaffolds functionalized with vitronectin and fibronectin to facilitate epithelial or fibroblast cell attachment.^19,20^ Nano-structured biomaterials demand for materials that can replicate biological properties, ensure biodegradability, and possess adjustable mechanical characteristics. While MPL with protein-based photoresists addresses these needs to some extent, the limited stability of the resulting 3D structures continues to pose a notable challenge.^21^

In this work, we demonstrate the capabilities of 3D MPL-nanolithography for structuring of new classes of amino acid-based phosphorodiamidate (APdA) monomers.^22,23^ For the experiments, three vinyl functionalized APdA monomers—Val-APdA-VE, Val-APdA-VC, and Ala-APdA-VC—were used as photoresists, with an additional 2 wt% of IC2959 photoinitiator. The APdA photoresist and writing parameters were optimized for MPL to achieve minimal lateral and axial feature sizes, and its 3D microstructuring capabilities were evaluated. To demonstrate the advantages of MPL over other optical lithography techniques, such as dynamic light processing (DLP) or nanoimprint lithography, which commonly require post-polymerization curing, we investigated the surface mechanical properties of MPL-fabricated APdA structures before and after UV curing. UV curing enhances polymer surface hardness, potentially affecting mechanotransduction and cell response^24^ or stress/wear-resistant surfaces.^25^ We quantified the mechanical properties of the polymeric structures under both wet and dry conditions, showing significant changes in the Young’s modulus without swelling of the structures. To offer options for surface functionalization that promote desired biological interactions with the APdA structures, we introduce a biotin-based surface modification strategy, allowing for versatile protein coating. We demonstrate its effectiveness through the immobilization of extracellular vesicles.

To quantitatively analyze the MPL writing performance of the photoresists, we designed an experiment that enables the simultaneous quantification of axial and lateral feature sizes of lines written in 3D, as shown in Figure 1a.^9,21^ Lines were structured perpendicular to the supporting structures. The three supporting structures out of Ormocomp^®^ were 5 µm in height and 100 µm in length with a spacing of 50 µm and 3 µm. Upon development, the short segment of the APdA line, positioned between the two narrow-spaced supporting structures (right), remains suspended, enabling the measurement of the lateral feature size. In contrast, the part of the APdA line written between the more widely spaced supporting lines tilts by 90° after washing, revealing the corresponding axial dimension of the same line (left). Figure 1b illustrates the three APdA monomers, each derived from valine- and alanine-based amino acids, which contain two vinyl groups used to formulate the three photoresists that are crosslinked by the Norrish Type I photoinitiator IC2959. In the formulations, 2 wt% IC2959 was directly mixed into the APdA monomers, and Val-APdA-VE was additionally heated to 40°C to decrease viscosity during the mixing process. For writing, a 20 µL droplet of photoresist was drop-cast onto a coverslip and excited in an inverted configuration using a 515 nm excitation wavelength (>290 fs pulse duration), focused through a 63x objective lens (NA = 1.4). For development, the uncured photoresist was removed by rinsing with 99% ethanol. The short monomer backbone and vinyl homofunctionality enable efficient crosslinking, allowing the fabrication of features at subdiffractional limits. Tilted lines have also been employed to quantify Young’s modulus using nanoindentation via atomic force microscopy (Figure 1c).

**Figure 1.**
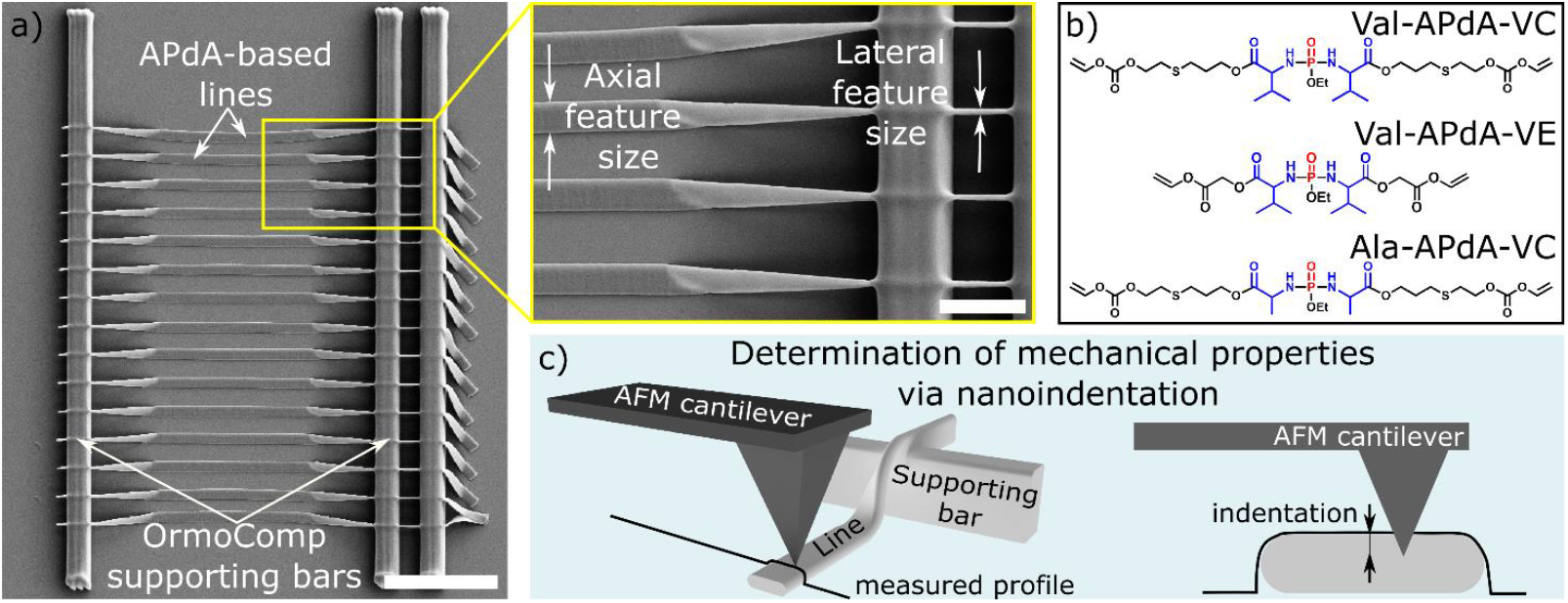
a) SEM image of the experimental setup used for characterizing the MLP writing performance. The long vertical bars represent the Ormocomp® support structures. APdA-based lines were structured perpendicularly to the supporting structures. Scale bar: 20 µm The short segment of the APdA-line, spanning between the two narrow supporting structures (see inset), remains suspended, allowing for the measurement of lateral feature size. The rest of the APdA-line written between the more widely spaced supporting lines gets tilted by 90° after washing, revealing the corresponding axial dimension of the same written line (see inset). Scale bar of inset: 5 µm b) Structural formulas of Val-APdA-VC, Val-APdA-VE and Ala-APdA-VC. Photoresists composed of each monomer and 2% IC2959 Norrish Type I photoinitiator were tested. c) Schematic representation of nanoindentation experiments on written APdA lines. c.) The sketch depicts the modalities of atomic force microscopy analysis for tilted polymeric lines.

Using this setup, we first quantified the spatial 2D writing properties by structuring lines on a glass coverslip, as well as creating 3D “hanging” lines and grid structures (Figure 2). 2D lines were structured in test arrays on standard glass coverslips (∼150 µm thickness) without any surface modification for a better relative comparison between the writing properties of the photoresists (see Figure 2a). For the analysis, we used the smallest reproducible lines, structured at 0.03 mm/s with 0.15 TW/cm^2^ and 100% reproducibility at the specified dosage. Thus, the average lateral feature sizes of 255 ± 20 nm, 140 ± 5 nm, and 250 ± 13 nm were determined for the 2D Val-APdA-VC, Val-APdA-VE, and Ala-APdA-VC lines (n=9 for each photoresist, exemplary lines shown in Figure 2a). Carbonate-based resins (VC) on glass showed diffraction-limited feature sizes ∼λ/2, while vinyl ester (VE) based monomers averaged ∼λ/3.7. The feature size may also be slightly increased due to the thickness of the metal coating (nominally 10 nm, required for SEM imaging), as each monomer absorbs the metal differently, as reflected in the contrast of the image. In comparison, the analysis of the hanging lines revealed overall smaller lateral feature sizes of 104 ± 11 nm, 83 ± 3 nm, and 136 ± 20 nm (averaged over n=9 lines, with exemplary lines shown in Figure 2b, top).

**Figure 2.**
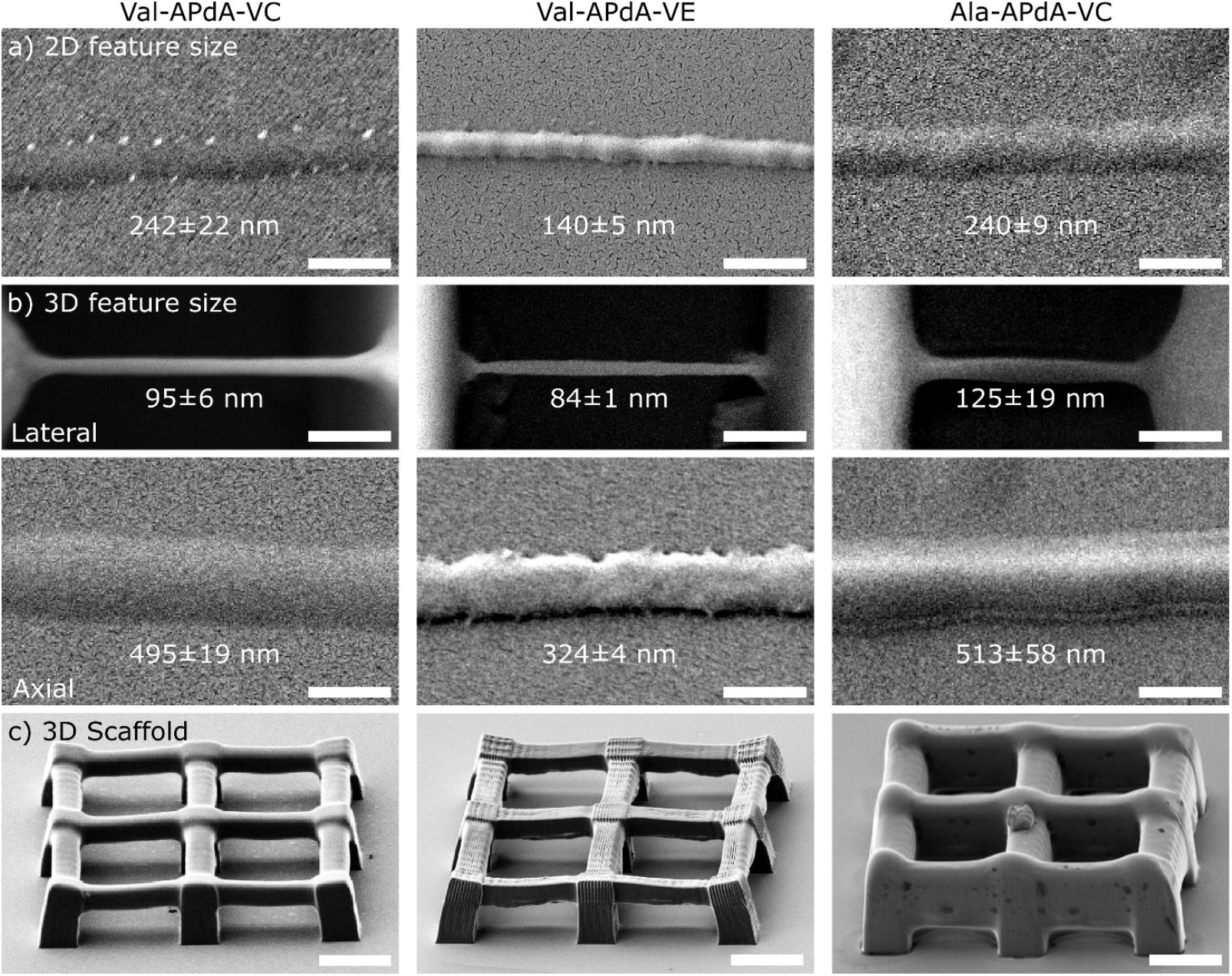
SEM images of 2D and 3D structures demonstrating the MPL performance of the resists. a) shows representative SEM images of the smallest, reproducible 2D lines made from Val-APdA-VC, Val-APdA-VE and Ala-APdA-VC (left to right) that have been developed. Lateral sizes of 242 ± 22 nm, 140 ± 5 nm, and 240 ± 9 nm have been determined for the lines presented. b.) SEM images of the 3D representative ‘hanging’- and 90° tilted-lines showing the lateral and axial feature sizes for all three photoresists. For the Val-APdA- VC, Val-APdA-VE, and Ala-APdA-VC lines, feature sizes of 95 ± 6 nm, 84 ± 1 nm, and 125 ± 19 nm, and axial feature sizes of 495 ± 19 nm, 324 ± 4 nm, and 513 ± 58 nm were determined, respectively. Scalebars in a and b are 500nm. The panels show the mean and standard deviation for the line measured at three points. c.) SEM images of 3D grid macro-structures (45×45×10 µm^3^) made of Val- APdA-VC, Val-APdA-VE, and Ala-APdA-VC monomers. Double exposure was used for Val-APdA-VC and Ala-APdA-VC, while quadruple exposure was used for Val-APdA-VE to achieve stable reproducible structures. Scale bars: 10 µm.

Nevertheless, the trend remains the same: VE exhibit the smallest lateral sizes (∼λ/6.2), whereas VC are generally larger (∼λ/4.3). Based on the results, we conclude that the slightly larger feature sizes observed in the Val-APdA-VC and Ala-APdA-VC 2D structures, compared to the Val-APdA-VE 2D structures, are likely attributable to the higher post-polymerization shrinkage and stronger adhesion of Val-APdA-VE compared to others. This enhances the structures’ resistance to washing, enabling the achievement of smaller feature sizes. The average axial feature sizes of the smallest tilted, hanging lines for Val-APdA-VC, Val-APdA-VE, and Ala-APdA-VC are 493 ± 30 nm, 355 ± 22 nm, and 519 ± 38 nm, respectively, enabling nanoindentation studies. The axial feature sizes in the tilted images are diffraction-limited, ∼ 440 nm (515 nm wavelength, NA = 1.4). In comparison, axial feature sizes for Val-APdA-VE lines measured at a 60° tilt for the hanging lines spanning the short-distance supporting bars were 280 ± 19 nm (see Supplementary Figure SI1), indicating that there is an adhesion between glass surface and lines causes the line broadening by up to 25%. However, these lines did not survive development along their full length and without corresponding tilted lines, thus, the respective Young’s moduli were excluded from statistics.

To show the capability for 3D structuring, we have written micrometer size 3D grid structures (Figure 2c, suppl. Figure SI2.). The pillar size was 5×5×10 µm^3^, and the bars in between were 5×3 µm^2^ wide/high and 15 µm long. The lateral line-to-line distance (hatching) was 500 nm, with an axial line-to-line distance (slice) of 750 nm. All lines were structured at 0.35 mm/s and 0.20 TW/cm^2^, with the VC-based photoresist illuminated twice and the VE-based photoresist illuminated four times to ensure mechanical stability during development, induced by direct rinsing of the structures with ethanol. Here, we exploit the properties of a ‘non-forgetting’ photoresist, which ‘remembers’ each exposure through induced crosslinking, resulting in increased viscosity. Thus, multiple exposures reduces the impact of residual single-photon excitation (residual absorption) in favor of the photoinitiator’s higher-order absorption of the APdA-monomers (absorption of monomers see in suppl. Figure SI3). The residual absorption contributes to heating, ionization and, consequently, micro-explosion of the photoresist.^26^ Reducing the MPL excitation decreases the excitation rate of photoinitiators per illumination cycle, resulting in lower monomer crosslinking and reduction in photoinitiator diffusion due to polymer gelation. In the second illumination step, a ‘post-curing’ process is introduced, where the more spatially confined photoinitiators within a higher-viscosity gel-phase polymer are subjected to double exposure, effectively reinitiating the polymerization reaction for the unreacted photoinitiators. This promotes mechanical stability; however, changes in viscosity and density (post-polymerization shrinkage) of the crosslinked polymer impact feature size and consequently structure shape. This is evident in the case of the 3D macro-structures, where the best results in writing precision were achieved for the most viscous valine-based resists. Ala-APdA-VC exhibited the largest deviations from the adjusted writing parameters, with the lateral bar height differing by nearly a factor of three.

Given their biodegradability and biocompatibility, APdA monomers have potential as biomaterials for tissue regeneration applications.^23^ Consequently, the mechanical properties of the hydrophobic APdA monomers under aqueous conditions are of significant interest. To investigate this, we employed AFM nanoindentation to analyze the swelling behavior and determine the Young’s modulus of MPL-structured APdA lines. We focused on quantifying the Young’s modulus of the MPL structure and characterizing the mechanical properties, systematically comparing this Young’s modulus with that of lines post-cured using UV light, a method commonly employed in optical 3D lithography techniques such as stereolithography (SLA), nanoimprint, or MPL post-curing. Tissue engineering applications involving UV hardening influences mechanotransduction between the scaffold and cells.^24^

First, we analyzed the surface mechanical properties of the monomers, noting that the photon density gradient of the MPL excitation profile leads to varying crosslinking efficiencies. The green voxel (Figure 3a) marks the polymerization profile above the threshold, reflecting the monomer’s crosslinking efficiency. It indicates a gradient in polymerization efficiency, correlating with the photon density-dependent energy dose. This leads to a gelation phase in the outer regions. Figure 3a shows the AFM topography and Young’s modulus map of the MPL-structured Val-APdA-VC line, while Figure 3b displays the AFM topography and Young’s modulus map of a Val-APdA-VC line UV-cured in a post-polymerization step. Measured Young’s modulus profiles of lines structured using MPL (Figure 3c) reveal a softer shell (18 MPa at minimum) around the hardened core (50 MPa). After UV curing, the 317± 30 nm FWHM broad softer shell hardens, resulting in a homogeneous Young’s modulus (48 MPa) profile across the line. The results prove that sufficient photoinitiators are present near the surface to restart the polymerization. It is important to note that the width of the measured soft shell depends on the properties of the cantilever and the tip used. The core-shell structure of the MPL features could be advantageous for cell-structure interactions, as the mechanical properties of local environments, such as matrix stiffness, are known to influence cell behaviors like migration, proliferation, differentiation, and metabolism.^24^

**Figure 3.**
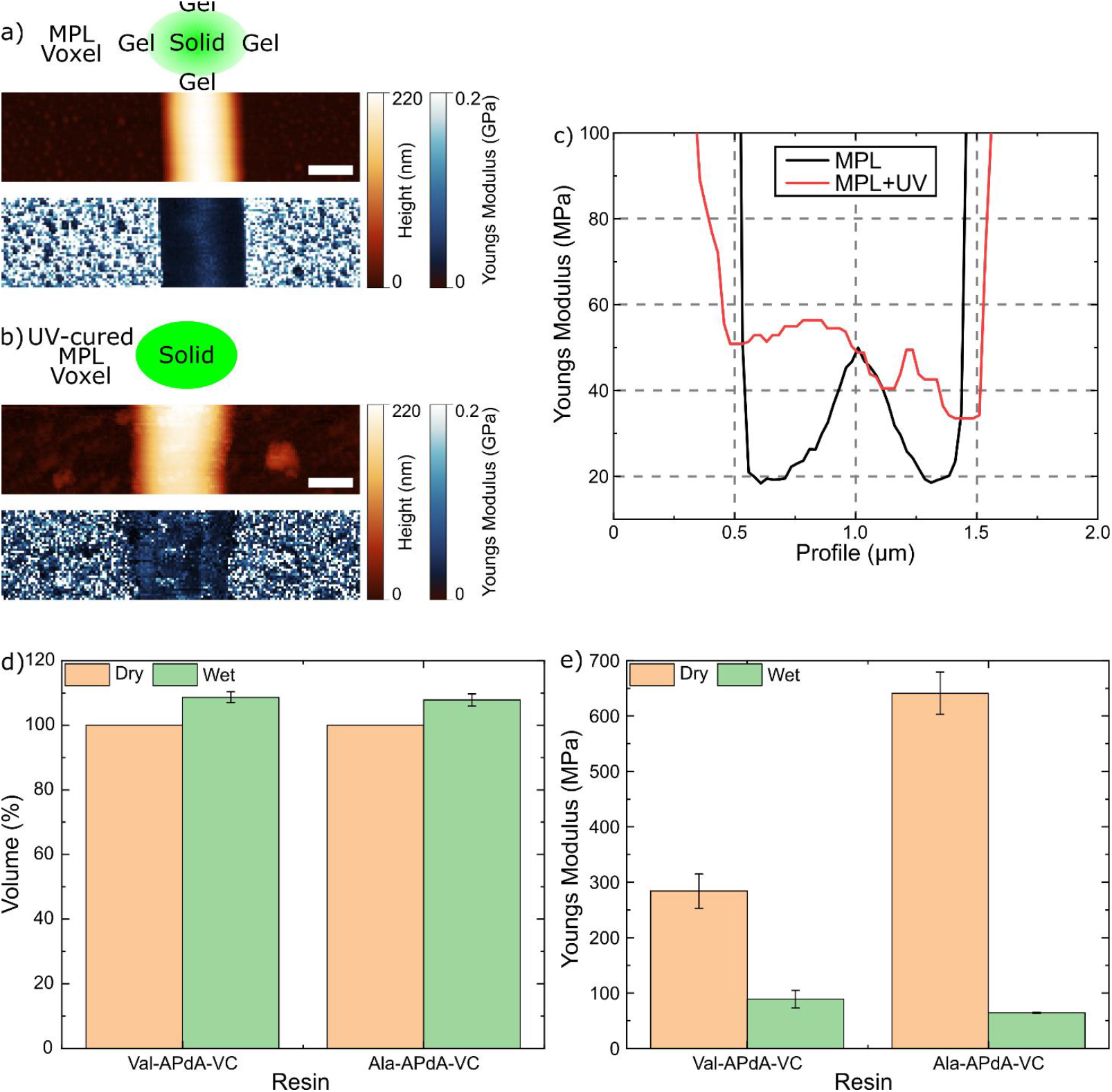
Mechanical properties of MPL-structured APdA lines. a) shows the topography of MPL-structured, Val-APdA-VC based line with the corresponding Youngs Modulus map. The green voxel depicts the polymerization profile above the MPL line threshold, showing a gradient in polymerization efficiency correlated with photon density that decreases from the inside outward. b) topography and a Young’s modulus map of a Val-APdA-VC line that was UV cured in a post-polymerization process (the same line as shown in a)). In contrast to a) the softer shell of the structure has been hardened by the UV-light under dry conditions. The results indicate that sufficient photoinitiations are present in the surface proximity to restart the polymerization. Scale bars in a) and b): 500 nm c) shows the Youngs Modulus profiles of Val-APdA-VC line, an MPL-structured (black) one and MPL-structured which has been UV-post cured (red). The Young’s Modulus analysis indicates that UV curing significantly hardened the MPL line’s surface, raising its Young’s Modulus to the maximum measured value and transforming the gradient Young’s Modulus of the MPL voxel into a more homogeneous material. d) diagram depicts the change in dimensions of Val-APdA-VC and Ala-APdA-VC structures under wet and dry conditions. The results show that the size of Valine- and Alanine-based structures increases by 8.6% and 7.8% respectively, measured by their height (integrated over the line width), when transitioning from dry to wet conditions. e) diagram showing the average Youngs Modulus measured for Val-APdA-VC and Ala-APdA-VC structures under wet and dry conditions. For the Valine based lines the Youngs Modulus (YM is the average over the FWHM width of the line) decreases from 284 ± 31 MPa (dry) to 89 ± 16 MPa (wet) when exposed to water. In case of Alanine-based monomers the change from 641 ± 38 MPa (dry) to 64 ± 1 MPa (wet) has been measured.

Next, we quantified and established a correlation between the structural size and Young’s Modulus for Val-APdA-VC and Ala-APdA-VC based structures, under wet and dry conditions (wetting: ∼15 hours). Owing to the relatively hydrophobic properties of the amino acids, significant swelling of the structure was not observed. The average swelling, measured by AFM, showed a total volume increase of 7.8% for Val-APdA-VC and 8.6% for Ala-APdA-VC, quantified based on the height of the structure. In contrast, the Youngs modulus changes significantly by 3.2 fold (from 284 ± 30 MPa (dry) to 89 ± 16 MPa (wet)) for Val-APdA-VC based polymer and 10 fold (from 641 ± 38 MPa (dry) to 64 ± 1 MPa (wet)) for Ala-APdA-VC based polymer ((YM is the average over the FWHM width of the line, 25 line profiles from each line and 5 technical replicas were analyzed). We also studied the dynamics of YM change during wetting. For this purpose, topography and YM were measured every 4 minutes (limited by AFM image acquisition time). We observed (see Supplementary Figure 4) a significant change in YM (195 MPa and 447 MPa for valine- and alanine-based resins, respectively) within the first 4 minutes. This suggests that the softening of nanostructures is not related to the degradation of monomers, as the determined degradation kinetics spanned tens of days.^23^ The change in YM due to swelling can also be largely excluded, as the change in the height of the structures was marginal (∼8% for both resins) compared to the observed change in YM.

**Figure 4.**
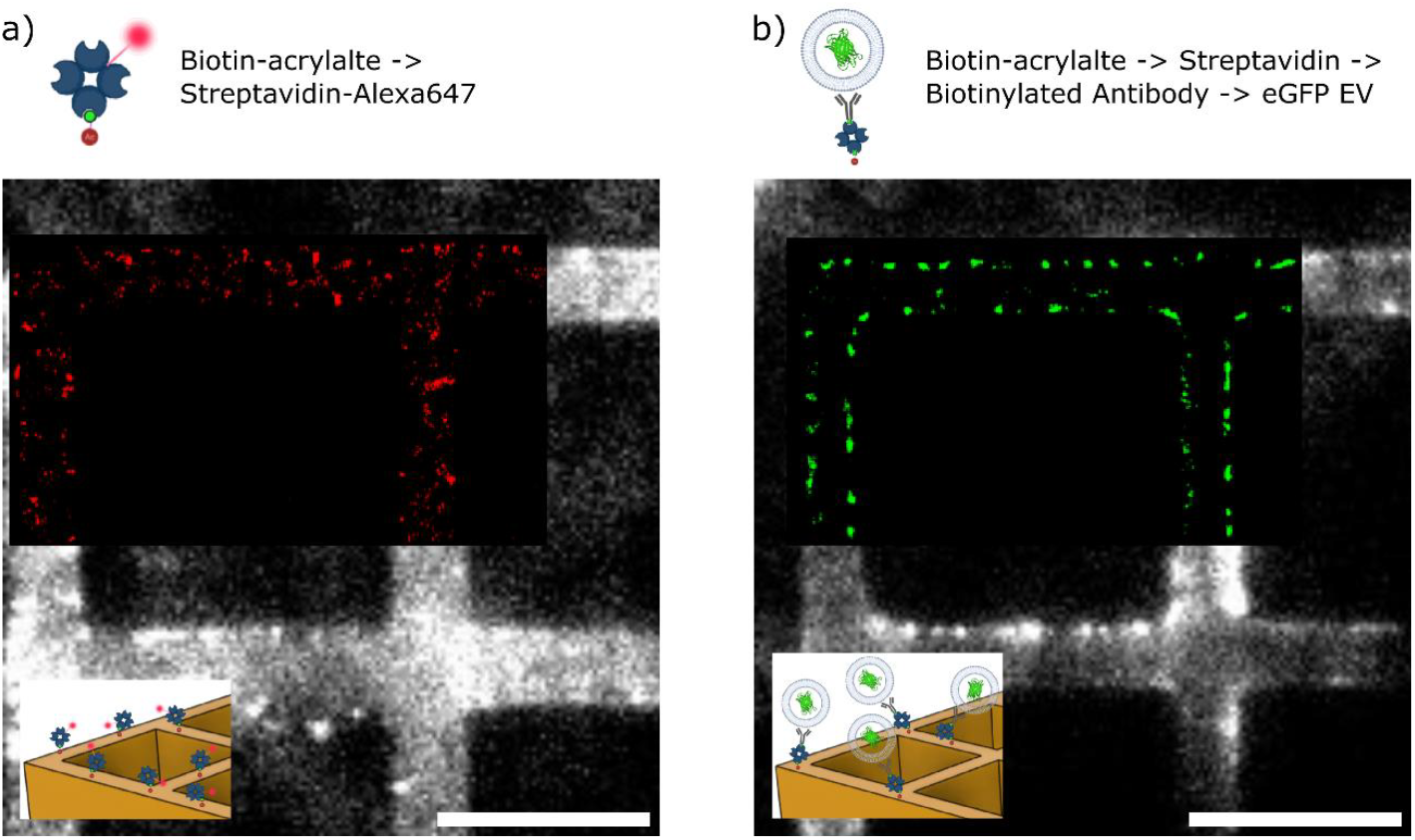
a) Fluorescence image of 3D biofunctional scaffold (made of Val-APdA-VC+2wt% of Biotin-acrylate) incubated for 1 min with 1 nM of Streptavidin-Alexa647. b) the same scaffold after subsequent incubation of biotinylated antibody (1 µg/mL anti-CD81 for 1 h) and eGFP-EVs (10^8^ particle/ml for 30 min).

A key focus was to analyze the effect of double exposure on the Young’s modulus of the structures, aiming to determine whether crosslinking efficiency is significantly affected, particularly in the context of its application to 3D structuring of macroscopic objects. We structured two lines on top of each other with an overlapping excitation volume (see suppl. Figure SI4). Nanoindentation measurements of Young’s modulus show that the solid core mechanical properties of the double-illuminated and single-illuminated lines are similar. As in all MPL-structured lines, the edges exhibit a softer shell. The measured Young’s modulus in the double-illuminated region was 57 ± 2 MPa (similar to single exposure), while lower values of 22 MPa were observed at the edges (FWHM=227 ± 22 nm). The results suggest that multiple exposures used for 3D structuring do not significantly affect the mechanical properties of the structures. However, they likely provide stabilization by compensating for post-polymerization shrinkage, which influences the shape and stability of the lines and is especially relevant for 3D scaffold fabrication. Interestingly, the variation in Young’s modulus does not correlate with the change in feature size during wetting.

Alongside biocompatibility, degradability and mechanical properties, surface functionalization is also of high relevance for the 3D structured materials for biological applications. We employed biotinylated methacrylates^27^ to functionalize the surface of the MPL structures, thus biotinylated methacrylates were added to the Val-APdA-VC photoresist (2 wt% Biotin-acrylate). The biotinylated methacrylates were evenly distributed within the photoresist, ensuring that a portion remained accessible for streptavidin binding on the surface of the polymerized material.

In order to quantify the specific binding a background (unspecific binding) needs to be determined. We compared the non-specific binding results of Streptavidin labeled with Alexa647N (Streptavidin-Alexa647N) to both biotinylated Val-APdA-VC and Val-APdA-VC structures by analyzing the specific and non-specific interactions at two time points. After 1 minute of incubation, the average nonspecific fluorescence increased by 127 ± 5 cnts. relative to the structures’ autofluorescence of 147 ± 5 cnts., while for the biotinylated structures, a twofold increase in fluorescence was observed, reaching 227 ± 4 cnts. For a more detailed analysis of the molecular distribution, single-molecule fluorescence microscopy was used to super-resolve the positions of the streptavidin on the surface of the MPL structures. Figure 4a shows an overlay of the fluorescence image with an image depicting the fluorescently localized Alexa 647-streptavidins on the structure, revealing a relatively homogeneous distribution of the proteins on the surface (estimated 40 streptavidin/µm^2^). After 20 minutes of incubation, the signal intensity for the unbiotinylated Val-APdA-VC was 655 ± 109 cnts. (scaffold autofluorescence: 227 ± 15 cnts), compared to 948 ± 90 cnts (scaffold autofluorescence: 235 ± 15 cnts). for the biotinylated Val-APdA-VC. The results validate the biotin surface modification of the APdA structures, distinguishing it from nonspecific protein binding (Supplementary Figure 6).

To explore the potential relevance of this coating for biological applications, streptavidin was utilized as an anchor for biotinylated anti-CD81 antibodies. CD81 (tetraspanin-28) is a transmembrane protein present in cell membranes as well as in extracellular vesicles. It is widely recognized as a marker molecule found in a significant proportion of mammalian extracellular vesicles produced by cells.^21,28–30^ To confirm the presence of antibodies on the surface, the structures were incubated with engineered extracellular vesicles (EVs) carrying a GFP-labeled CD63 transmembrane protein derived from HEK-cells, as described in ^31,32^. The fluorescence images display the GFP signals of surface-bound EVs, localized at the single-particle level. Figure 4b depicts the average EV fluorescence signal and their localized positions on the surfaces of the 3D structures. Specific EV binding to streptavidin-modified structures showed an average increase of 43 ± 2 cnts. over non-specific binding (estimated 17 EVs/µm^2^). Supplementary Figure 7 includes additional fluorescence and super-resolution images taken at the cross-section of the structure, confirming that streptavidin functionalization extends along the sidewalls of the 3D scaffold. The results confirm that biotinylated structures enable streptavidin binding and antibody immobilization, supporting target molecule binding (e.g., EVs) and demonstrating suitability for two-color, single molecule fluorescence microscopy.

In conclusion, we have demonstrated the potential of 3D multiphoton lithography (MPL) for the precise structuring of three new amino acid-based phosphorodiamidate (APdA) monomers. By optimizing the APdA photoresist formulations and MPL writing parameters, we achieved sub diffraction-limited feature sizes for both 2D and 3D structuring. In general ester (VE) based monomers resulted in smaller feature sizes (down to 140 nm in 2D and 84 nm in 3D) compared to carbonate-based (VC) resins (down to 240 nm in 2D and 95 nm in 3D)). We further evaluated the mechanical properties of MPL-fabricated APdA structures under wet and dry conditions, revealing significant decrease of Young’s moduli (from 3 to 10 fold, depending on composition) without notable swelling (∼8%). Additionally we showed a biotin-based surface functionalization strategy that enables the immobilization of extracellular vesicles, showcasing the potential for tailored biological interactions. This versatile approach opens pathways for designing advanced biomaterials for applications in tissue engineering, drug delivery, and biointerface engineering.

## Associated Content

### Supporting Information

The Supporting Information is available free of charge. Additional characterizations, including SEM images, dynamics of YM change, YM for double illuminated lines, fluorescence image, and details of the experimental methods (PDF)

## Author Information

Corresponding Author

Dmitry Sivun – School of Medical Engineering and Applied Social Science, University of Applied Sciences Upper Austria, 4020 Linz, Austria

## Acknowledgments

This work was supported by the Austrian Research Promotion Agency (FFG) project FFG898921 “Nano-Carriers”. We would like to thank Heidi Piglmayer-Brezina for taking the SEM images, Ilse Kammerhofer for support with the administration organization.

