## Supplemental Information for "3D Multiphoton Nanolithography with Bioresorbable Amino Acid-Based Resins"

for

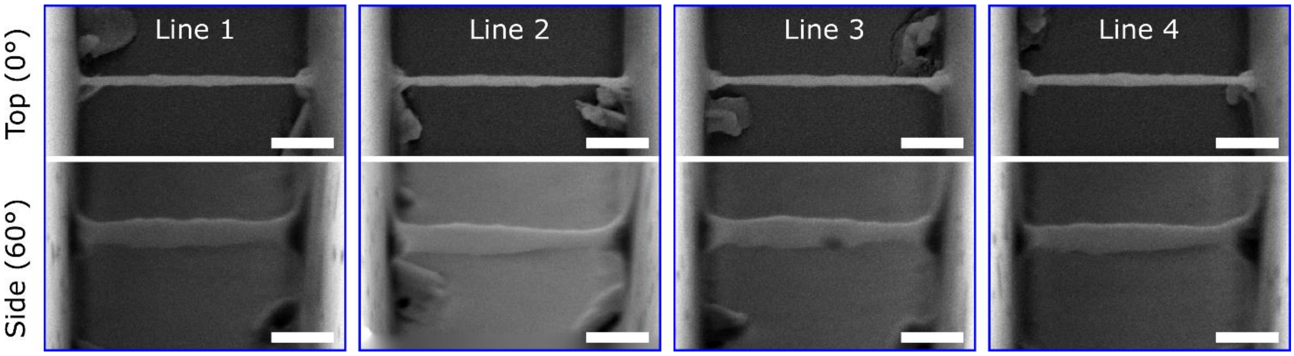


Supplementary figure 1. SEM images of lateral (upper panel) and axial (lower panel) features size for 4 technical replicas of Val-APdA-VE “hangigng” 3D line (as presented in fig 2b middle column). The axial images are taken at 60° sample tilt. Scale bars: 500 nm.

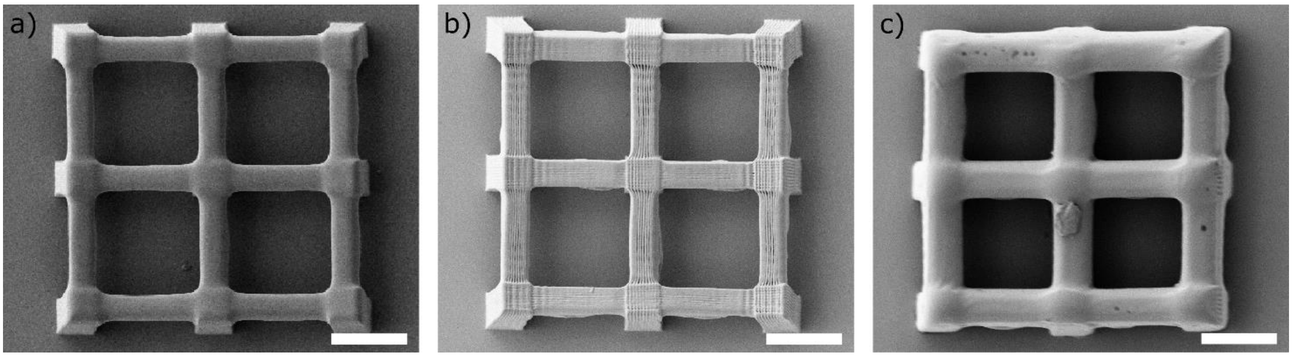


Supplementary figure 2.: The images show top SEM images of the 3D structures represented in figure 2c. The images confirm the claim that Val-APdA-VE structures deliver the best performance in terms of detail (feature size), even after multiple exposures. Scale bars: 10 µm.

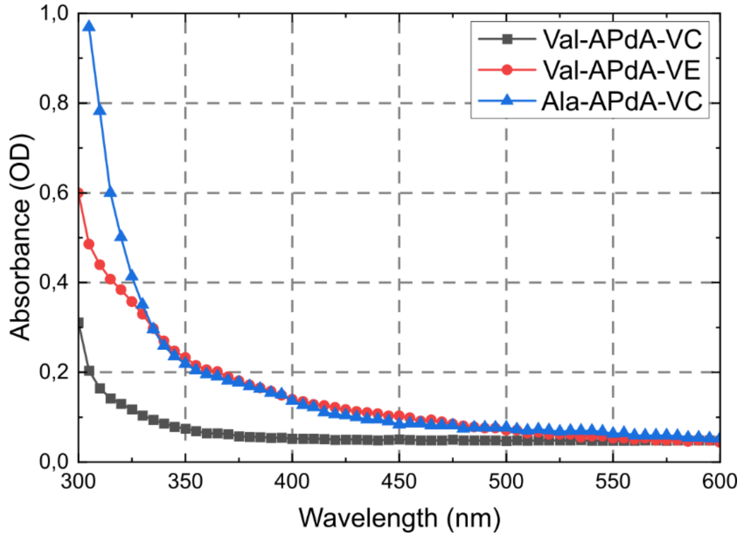


Supplementary figure 3.: Absorption spectra of three used monomers.

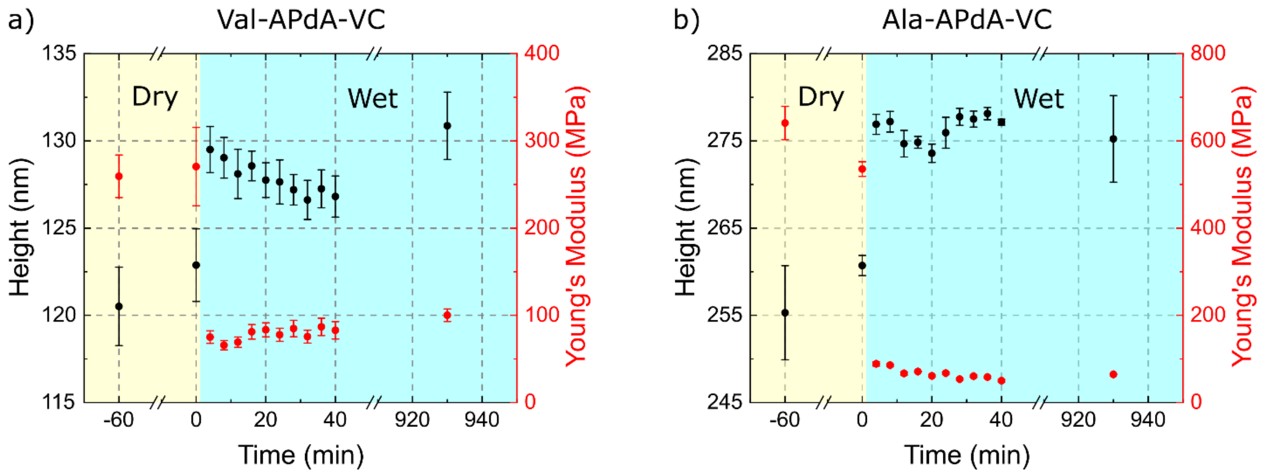


Supplementary figure 4.: The diagrams display the time dependent change in line dimensions and Youngs Modulus upon wetting. The data reveals a fast wetting within the first 7 min with a corresponding change in line dimensions and Youngs Modulus. After wetting, both parameters do not change significanlty for 930 minutes.

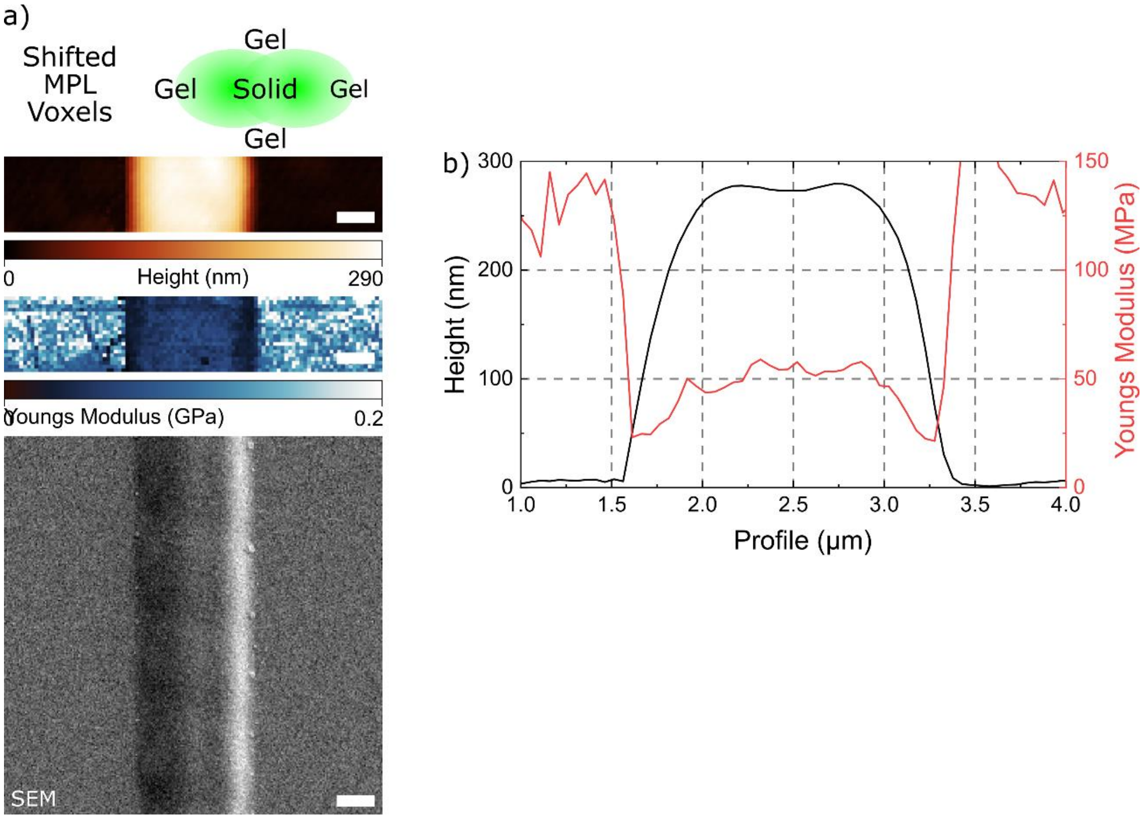


Supplementary figure 5. Line written with a 500 nm axial shift of MPL focus. a) shows (from right to left) the schematics, AFM topography and Young's modulus, and SEM image of the line. b) cross-sections of the topography and Young's modulus presented in a). Cross-section profile is averaged over 1µm. Scale bars: 500 nm

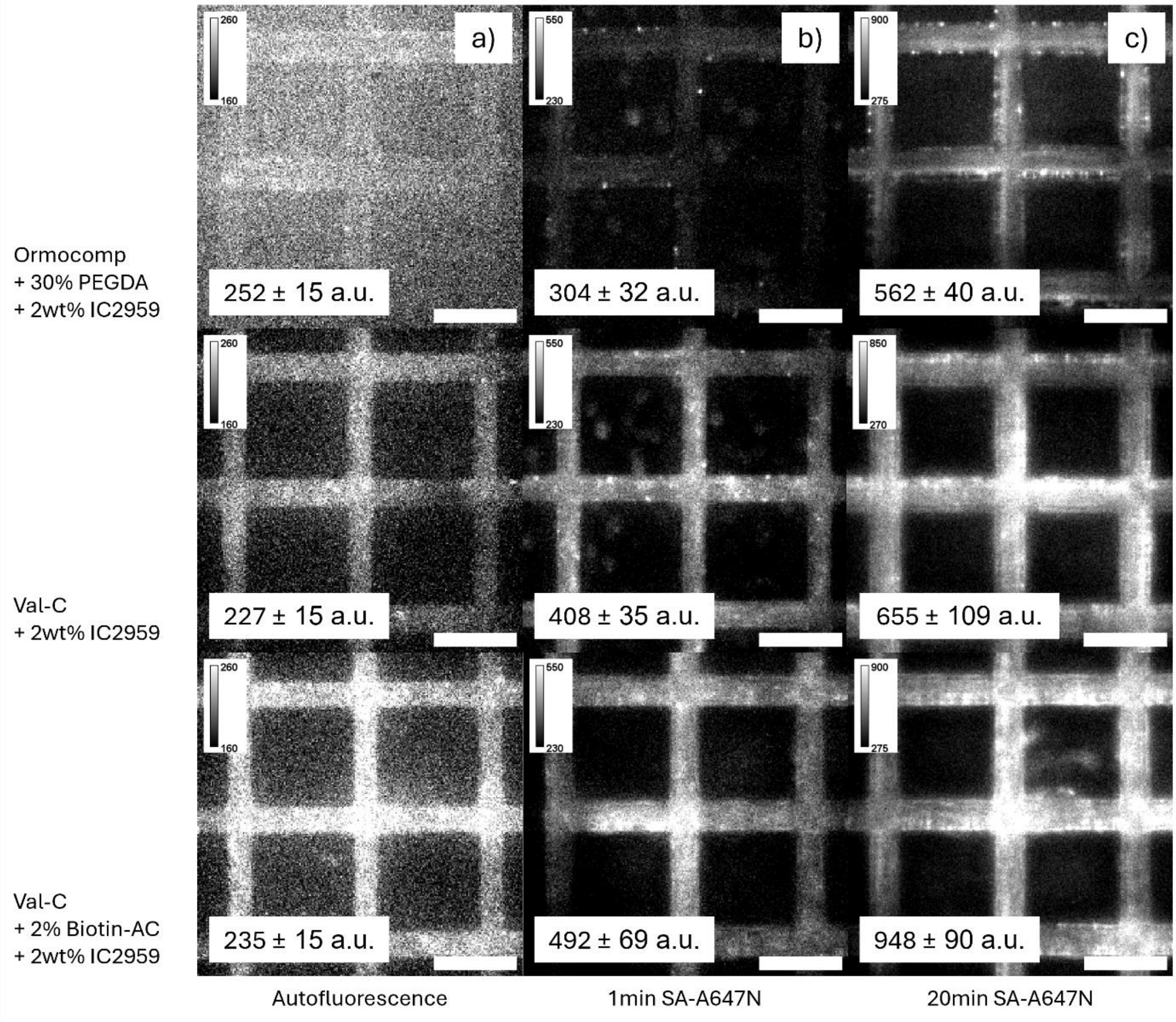


Supplementary Figure 6. Fluorescence images of specific and unspecific binding of Streptavidin-Alexa647N to three photoresist compositions. Ormocomp® + 30% poly (ethylene glycol) diacrylate (PEGDA) was used as a protein-repellent control. Unmodified Val-C was used for quantification of unspecific binding and Val-C with Biotin-AC was used for specific binding of streptavidin. Collums a) b) and c) show the autofluorescence, the fluorescence signal after 1 min incubation of Streptavidin-Alexa647N and the fluorescence signal after 20 min incubation with Streptavidin-Alexa647N for the three photoresist compositions, respectively. Scale bars 10 µm.

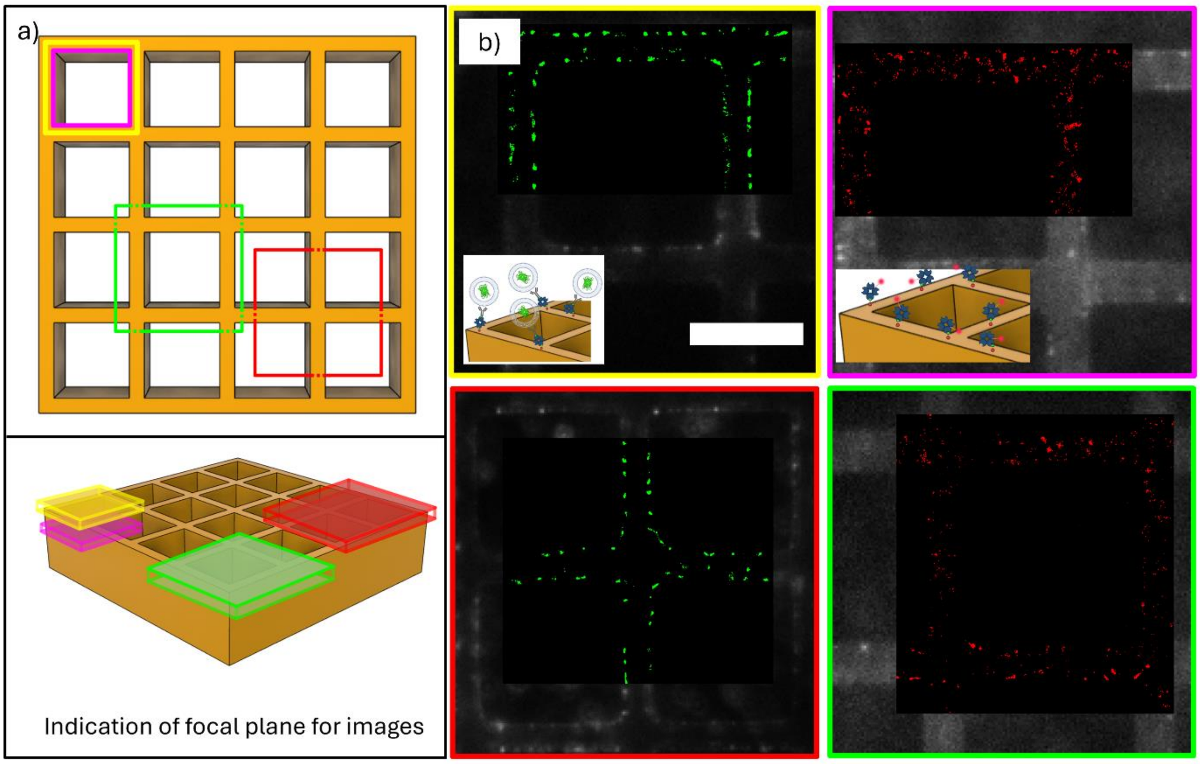


Supplementary Figure 7. Fluorescence and super-resolution images of surface immobilized streptavidin and eGFP-EVs. a) shows a schematic of the focal planes on the 3D structure where the images were taken. b) fluorescence images of eGFP-EVs (left) and Streptavidin-Alexa647N (right) with super-resolved overlays of the focal planes indicated in a). Scale bar 10 µm for all images.

### Materials and Methods

*Materials*

Details on the synthesis of APdA monomers are available in our previous publication.[1]

*Multi-Photon Lithography (MPL)*

All presented structures were fabricated with a customized multiphoton lithography (MPL) system from Workshop of Photonics (WOP, Lithuania). The setup featured an fs-pulsed laser (CARBIDE, 515 nm, 1 MHz repetition rate, 290 fs pulse duration, Light Conversion, Lithuania). The stated peak intensities were calculated as focal plane values within a diffraction-limited excitation volume, based on the average power measured at the objective lens input, accounting for its transmission. Excitation laser was focused to the sample plane by 63x oil immersion objective lens (NA=1.4, Zeiss) and writing is performed by moving sample via 3-axis stage (AEROTECH Nanopositioner, USA). Post-fabrication development involved rinsing the samples with 99% ethanol to remove uncured resin.

Atomic Force Microscopy (AFM)

Young's modulus was measured using an atomic force microscope (JPK NanoWizard 4, Germany) operated in QI™ mode. The AFM was integrated with a Zeiss AxioObserver inverted epifluorescence microscope. The measurements utilized a PPP-FMR cantilever (Nanosensors, Germany) with a nominal spring constant of 2.8 N/m. Prior to each measurement, the cantilever's spring constant and sensitivity were calibrated on the substrate near the polymer lines using the contact-based calibration method (JPK, Germany).

Scanning Electron Microscopy (SEM)

SEM images were obtained using a Zeiss 1540XB SEM after evaporating approx. 10 nm of gold. SEM of three-dimensional structures was performed at normal incidence (0°) and by tilting the samples by 60°.

Single Molecule Fluorescence Microscopy (SMFM)

Images were acquired using a modified Olympus IX81 inverted epifluorescence microscope with an oil-immersion objective lens (60x/ NA = 1.42, Zeiss) and an additional 1.6x magnification tube lens for a final magnification of 96x. The sample was positioned on a XYZ piezo stage (P-733.3DD, Physical Instruments, Germany) with nanometer precision on top of a mechanical stage with a range of 1 × 1 cm (JPK Instruments, Germany). The sample was illuminated in the green channel with a 488 nm laser (Toptica Photonics, Germany) and in the red channel with a 640 nm laser (Toptica, Photonics, Germany) in wide-field configuration. The signal was detected using an Andor iXonEM+ 897 (back-illuminated) EMCCD camera (16 µm pixel size). The following filter sets were used: dichroic filter (ZT405/488/561/640rpc, Chroma, Olching, Germany), emission filter: BP525/50 or HQ 700/75 M (both Chroma Technology GmbH, Germany) for the green and red channels respectively. The parameters for dSTORM imaging were as follows: $t_{ill}$ = 20 ms, $t_{delay}$ = 20 ms, $N_{accrued\ frames}$ = 500, $I_{640nm}$ = 2.91 kW/cm$^2$, $I_{488nm}$ = 3.40 kW/cm$^2$. The 3D localization analysis of Streptavidin-Alexa647N and eGFP-EVs was performed using custom–built software – 3D STORM tools.[2]

The results revealed a single molecule intensity of an Alexa647-streptavidin molecule: 389 ± 46 a.u. with a localization position accuracy of 48 ± 15 nm (SNR = 23.5). For the single EV fluorescence revealed a single EV intensity of 810 ± 54 a.u. (average degree of labelling = 3.2) with a localization position accuracy of 40 ± 12 nm (SNR = 16.7).

To minimize unspecific binding on the polymer structure a 2% solution of bovine serum albumin (Sigma Aldrich) in phosphate buffered saline (PBS) was incubated for 1 h. To reveal the binding capabilities of the Biotin-acrylate modified Val-APdA-VC 1 nM Streptavidin-Alexa647N (ThermoFisher) in PBS was applied for 1 min. After washing 15x with PBS, the sample was incubated with 1 µg/mL biotinylated antibody against CD81 (ThermoFisher) und subsequentially washed 20x with PBS. eGFP labelled EVs from HEK cells were captured by the antibodies ($10^8$ particles/mL for 30 min).

*Extracellular Vesicles*

Transfections, cell culture, and subsequent extracellular vesicle (EV) isolation were carried out following previously published protocols.[3] Briefly, HEK293T cells (ATCC #CRL-3216) were maintained in Dulbecco's modified Eagle's medium (DMEM, Thermo Fisher, Waltham, MA, USA), high glucose, supplemented with 10% fetal bovine serum (FBS, Thermo Fisher) and 1% penicillin/streptomycin (complete medium) at 37 °C in a 5% $CO_2$ atmosphere. For overexpression, cells were transfected with 1 µg of the GFP-CD63 plasmid (Addgene #62964) using Endofectin Max reagent (Genecopoeia, Rockwell, MD, USA) according to the manufacturer's instructions. Following several weeks of cultivation, GFP+ cells were sorted using fluorescence-activated cell sorting (FACS). After sorting OE-GFP-CD63 HEK293 cells were cultured as parental cells. To deplete EVs, a 20% FCS solution was centrifuged at 110,000×*g* for 12-h. For collection of the conditioned media, the media was changed to OptiMEM (Gibco, Carlsbad, CA, USA) containing 2% EV-depleted FCS, supernatants were harvested after 24 hours from approximately 130 × $10^6$ cells. To remove cellular debris, apoptotic bodies, and larger EVs, the supernatants underwent sequential centrifugation at 200×g (5 min), 2000×g (10 min), and 10,000×g (30 min) at 4 °C. Smaller EVs were isolated by ultracentrifugation at 110,000×g for 1 hour and 10 minutes at 4 °C using a Sorvall MX150 ultracentrifuge (Thermo Fisher Scientific, Waltham, MA, USA). The resulting EV pellet was washed with PBS, re-concentrated by ultracentrifugation, and resuspended in 30 µL of PBS. Samples were stored at −80 °C. The protocol was adapted from Bobrie et al.[4]
